# Task dependent cortico-cerebellar responses to delayed visual movement feedback

**DOI:** 10.1101/2025.04.17.649407

**Authors:** Gesche Vigh, Jakub Limanowski

## Abstract

Forward models in the brain issue predictions of sensory movement consequences that can establish self-other distinction through comparison with actual feedback. But forward models can also be updated by sensory prediction errors, as in sensorimotor adaptation. Disentangling the neuronal correlates of processes related to sensorimotor incongruence detection from those related to adaptation can be challenging. Here, we approached this challenge with a novel, virtual reality based hand-target tracking task that allowed us to manipulate the behavioral relevance of delayed visual movement feedback, while recording hemodynamic brain activity with functional magnetic resonance imaging (fMRI). Participants performed continuous right hand grasping movements to track a target oscillation with either their real, unseen hand or with a glove-controlled virtual hand. The movements of the virtual hand were delayed in a roving oddball fashion. Therefore, tracking with the virtual hand (VH task) required visuomotor adaptation, whereas tracking with the real hand (RH task) required ignoring visuomotor delays. VH > RH task execution produced stronger activity in the left posterior parietal cortex (PPC) and the bilateral extrastriate visual cortex. Delays correlated with activity in several regions, prominently including right temporoparietal regions, in both tasks. Crucially, the cerebellum showed a stronger delay dependent activation, and increased functional connectivity with the PPC, in the VH > RH task. Delay changes and errors correlated with activity in the anterior insulae (AI), and more strongly so in the VH>RH task. Thus, the instructed behavioral relevance of delayed visual movement feedback enhanced responses of the cerebellum and PPC, and their communication, likely for visuomotor adaptation. In contrast, temporoparietal regions compared predicted and actual visual movement feedback irrespective of its behavioral relevance, while the AI signaled visual mismatches particularly when these were behaviorally relevant.

## Introduction

Within computational architectures of motor control, forward models predict the sensory consequences of the to-be-executed muscle movements based on corollary signals from the motor system (Desmurget & Grafton, 2000; Parr et al., 2021; Shadmehr & Krakauer, 2008; Wolpert, Miall, et al., 1998). These predictions are essential for on-line motor control, but can also be used for distinguishing self-produced from externally generated sensations and, thus, for bodily self-other distinction (Blakemore et al., 1998; Wolpert & Flanagan, 2001; Jeannerod, 2004). Thus, in healthy people, sensory movement consequences in line with predictions by forward models are self-ascribed, whereas unpredicted sensations are perceived as having been externally caused (Gallagher, 2000; Farrer & Frith, 2002; Haggard, 2017; Synofzik, Vosgerau, et al., 2008). This is thought to be implemented by ‘comparator’ modules, in which the neural activity produced by predicted (‘self-generated’) sensations is relatively attenuated compared to that evoked by externally generated ones (Frith, 2000; Synofzik, Vosgerau, et al., 2008; Wolpert & Flanagan, 2001).

Following this logic, the experimental introduction of sensorimotor incongruence has been widely used to study the neuronal basis of comparator models for self-other distinction (see the reviews and meta-analyses by Brass et al., 2009; David et al., 2008; Seghezzi et al., 2019; Sperduti et al., 2011). Most common is the introduction of spatial or temporal incongruences to visual movement feedback, through mirrors, manipulated videos, or more recently, virtual reality. Moving under visuomotor incongruence has been associated with activity increases in (predominantly right) temporoparietal and inferior parietal regions, premotor cortices, the cerebellum, and the anterior insulae (Balslev, 2004; Balslev et al., 2006; Brass et al., 2009; Christensen et al., 2007; Farrer et al., 2008; Farrer & Frith, 2002; Kilteni & Ehrsson, 2024; Leube, 2003; Limanowski et al., 2018; Nahab et al., 2011; Ohata et al., 2020; Tsakiris et al., 2010; Uhlmann et al., 2020; Yomogida et al., 2010). The common interpretation of these results is that at least some of these regions compare predicted with actual visual movement feedback, and detect and signal mismatches—thus contributing to evaluating agency (see above). Human brain imaging studies have provided considerable evidence that regions around the right temporoparietal junction (including the angular gyrus (AG), supramarginal gyrus (SMG), and posterior superior temporal sulcus (pSTS)) play a key role in implementing such visuomotor comparisons (Balslev, 2004; Leube, 2003; Limanowski et al., 2018; Van Kemenade et al., 2017, 2019).

However, the brain’s motor system can learn novel visuomotor relationships; i.e., its forward models can be updated by prediction errors arising from unpredicted visual movement consequences. This is known as visuomotor adaptation. Notably, several of the above ‘delay-sensitive’ areas are also evidently involved in visuomotor adaptation; thereby, previous work has specifically implied the PPC and the cerebellum (Grafton et al., 2008; Küper et al., 2014; Limanowski et al., 2017; Ogawa et al., 2006, 2007; Synofzik, Lindner, et al., 2008; Tseng et al., 2007; Tzvi et al., 2017, 2020). Commonly, the PPC and cerebellum are thought to implement said forward models (Desmurget et al., 1999; Hartwigsen et al., 2012; Miall et al., 1993; Wolpert, Goodbody, et al., 1998; Wolpert, Miall, et al., 1998). In particular, updating the cerebellar forward models by sensory prediction errors seems to be a cornerstone of sensorimotor adaptation (Galea et al., 2011; Izawa et al., 2012; Block & Celnik, 2013; Chapman et al., 2010; Danckert et al., 2008; Luauté et al., 2009; Miall et al., 2001; Synofzik, Lindner, et al., 2008; see Tzvi et al., 2022, for a review). This means that cortical and cerebellar activations previously observed during movements under visuomotor incongruence could, in principle, indicate the detection or processing of visuomotor mismatches *and/or* processes related to feedback learning and adaptation. Disentangling these processes in a single study design is challenging (see Grafton et al., 2008; Limanowski et al., 2017; Ogawa et al., 2007, for previous attempts).

Here, we approached this challenge with a novel, virtual reality based hand-target tracking task that allowed us to manipulate the behavioral relevance of delayed visual movement feedback, while recording hemodynamic brain activity with fMRI. We have previously shown that instructed behavioral relevance can up- or down-regulate the processing of visual (vs proprioceptive) movement feedback (Limanowski, 2022; Limanowski & Friston, 2020). Here, we tested whether and how the instructed behavioral relevance of visual movement feedback affects responses in brain regions supposedly implementing forward models or comparator modules, respecitvely (i.e., such as the PPC, the cerebellum, or the temporoparietal cortex).

Participants performed continuous right hand grasping movements to track a virtual target oscillation with either their real, unseen hand (RH task) or with a virtual hand (VH task) that they controlled via a data glove (Fig. 1). The movements of the virtual hand were permanently delayed with respect to the real hand’s movements, while the amount of delay was repeatedly varied in a roving oddball fashion. Crucially, the VH task required repeated visuomotor adaptation and, consequently, attention to visuomotor delays and changes therein. Conversely, the RH task required ignoring the delays and changes in order to keep the real hand movements aligned with the target. This established the behavioral relevance of (delayed) visual movement feedback, under identical visuomotor delay sequences. With this task design, we aimed to disentangle the hemodynamic correlates of: visually guided action control (VH > RH task); visuomotor incongruence detection (effect of delay and changes in visuomotor mapping, which we expected in regions implementing comparator modules); and feedback learning and visuomotor adaptation (i.e., stronger delay-dependent responses in the VH > RH task, which we expected in regions implementing forward models).

**Figure 1:**
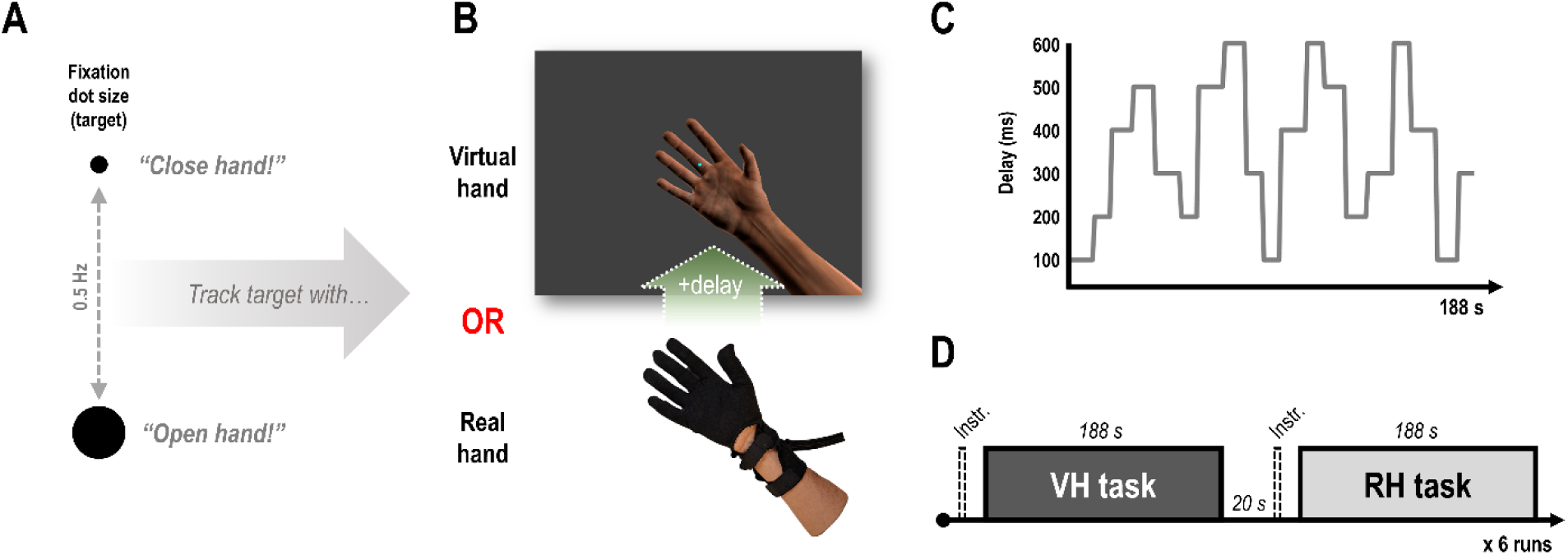
Hand-target tracking task. **A:** The participants’ task was to track the 0.5 Hz oscillatory size change of a central fixation dot with right-hand grasping (open-and-close) movements. **B:** Participants wore an MR compatible data glove on their right hand, with which they controlled a photorealistic virtual hand model. During the experiment, the virtual hand movements were permanently delayed with respect to the real hand movements. This implied that only one of the hands (virtual or real) could be synchronized with the target oscillation, while the other one would consequently move out of synchrony. For instance, if the real hand would be aligned with the target, the delayed virtual hand movements would lag behind it. Throughout the experiment, participants were instructed to track the target oscillation with either the virtual hand (VH) or their real, unseen hand (RH). This means they either had to adapt to the visuomotor delays (VH task) or ignore them (RH task). **C:** During the continuous movement task, the amount of visuomotor delay varied between 100 and 600 ms; i.e., changes in delay were introduced gradually every 4-6 movement cycles in a roving oddball fashion (requiring repeated adaptation in the VH task). The plot shows the delay sequence of an exemplary task block. **D:** Schematic trial structure of one of the 6 fMRI runs, which each participant completed. In each run, the same delay sequence (e.g., see panel C) was presented after an instruction to track the target with the virtual hand (VH task) and the real hand (RH task; the task order was randomized).

## Materials and Methods

### Participants

Twenty healthy, right-handed participants (15 female, mean age = 28.1 years, range= 22-38, normal or corrected-to-normal vision) took part in the experiment. The sample size was based on our previous studies using similar VR hand-target tracking tasks (Limanowski et al., 2017; Limanowski & Friston, 2020). Participants had successfully completed a training session prior to the fMRI experiment; i.e., only participants who showed a significant difference in real hand phase shift between the VH and RH task, indicating visuomotor adaptation vs no adaptation (see below), were invited to the fMRI experiment. The experiment was approved by the ethics committee of the Technische Universität Dresden and conducted in accordance with this approval.

### Experimental design and procedure

During the fMRI experiment, participants lay comfortably inside the scanner. On their right hand, they wore an MR-compatible data glove (5DT Data Glove 14 Ultra MRI, 60 Hz sampling rate, communication with the PC via USB 1.1), which measured each finger’s flexion via sewn-in optical sensors. The glove data were recorded and fed to a photorealistic right virtual hand model; i.e., participants were able to move the virtual fingers (which received an average of the glove’s sensor values to ensure smooth motion, cf. Limanowski & Friston, 2020). During the entire experiment, participants only saw the virtual hand (which they controlled by their real hand movements), but not their real hand (which was placed outside their field of view). The data glove was calibrated individually to ensure a good fit and coherent movement range across participants. Through a mirror on the head coil, participants saw the virtual environment (hand, target, and instructions) projected on a screen (projector model: Casio XJ-M156, Laser/LED hybrid, 1024 x 768 pixels resolution, 60 Hz refresh rate). The virtual environment was created within the Blender 3D graphics software package (https://www.blender.org, Version 2.79).

Participants were instructed to fixate a central dot in the middle of the screen, which changed size (∼12% size change) sinusoidally at 0.5 Hz (i.e., one full cycle every 2 s, Fig. 1A). Their task was to track this oscillatory size change with grasping (open-close-open) hand movements in continuous movement blocks of 188 s duration (94 grasping cycles, see Fig. 1C and below). As in previous studies (Limanowski & Friston, 2020), we used an oscillating fixation dot as the target to ensure that participants kept looking at the center of the screen in both conditions; we chose a central oscillation instead of e.g. a spatial (moving) target to eliminate potential eye movements resulting from smooth pursuit of the target as much as possible.

Throughout the experiment, the virtual hand movements were permanently delayed with respect to the actually executed (real hand) movements (Fig. 1B). Thus, the (seen) virtual and (unseen) real hand positions were always incongruent; meaning that only one of the hands’ (virtual or real) grasping movements could be synchronized with the target oscillation at a time—while the respective other hand would necessarily move out of synchrony.

Crucially, participants were instructed to track the target oscillation either with the virtual hand (VH task) or with their real, unseen hand (RH task). Thus, if participants wanted to track the target with the virtual hand (VH task), they had to phase-shift their real hand movements to compensate for the visual feedback delay; i.e., they had to perform visuomotor adaptation. Conversely, to maintain alignment of their unseen real hand movements with the target oscillation (RH task), participants needed to ignore the visuomotor delays. This effectively rendered visual movement feedback – and, importantly, visuomotor delays and their changes – behaviorally relevant (VH task) or irrelevant i.e. distractors (RH task).

During the continuous movement blocks, visuomotor delays varied between 6 different delay levels (100-600 ms in 100 ms increments) in a roving oddball fashion, in pseudorandomized sequences (cf. (Vigh & Limanowski, 2025): Each of the 6 delay levels was presented 3 times per run, for 4-6 movement cycles (8-12 s), before a different (i.e. shorter or longer) delay was introduced (18 delay changes per run, see Fig. 1C for an exemplary sequence). The delay changes were introduced gradually (in steps of 33 ms) to avoid sudden jumps in the visual feedback and to target implicit adaptation where possible (Limanowski et al., 2017; Tzvi et al., 2022). Each delay sequence consisted of 94 movement cycles (188 s). We kept the lengths of each delay period (max. 12 s) deliberately short (vs e.g. 30 s in Limanowski et al., 2017) to maximize the number of delay changes within the scanning time. Thus, our continuous movement task design was not aimed at trial-by-trial learning (and we did not expect participants to reach complete adaptation in the VH task), but optimized in terms of manipulating the cognitive-attentional set via the behavioral relevance of visuomotor delays and their changes; i.e., requiring constant vigilance and repeated re-adaptation in the VH task.

Each participant completed 6 scanning runs in total. In each run, the same 188 s long delay sequence was presented twice in consecutive movement blocks separated by 20 s rest periods; i.e., under the instruction to track the target with the virtual hand (VH task) and the real hand (RH task), see Figure 1D; the task order was randomized. With this, we ensured that the VH and RH conditions contained exactly the same delays and changes. Prior to each movement block, an instruction appeared on screen for 3 s (“virtual”/“real” in German). The text—and the fixation i.e. target dot in the subsequent movement block—were colored pink or turquoise (randomized across participants) to help participants remember the current instruction during movement. After each run, participants were able to rest until they were ready to continue.

### Behavioral data analysis

In order to quantify task compliance and compare adaptive responses between tasks (i.e., whether participants showed signs of visuomotor adaptation in the VH but not the RH task), we calculated the phase shift between the target oscillation (the phasic size change of the fixation dot) and the average hand movement (real hand) trajectory executed by each participant in each condition. Here, we excluded the first two movement cycles of each delay train, to eliminate effects of the gradually introduced delay changes; i.e., to focus on periods of the movements, in which visuomotor adaptation could be expected. Note that, to re-align the delayed virtual hand movements with the target oscillation (i.e., VH task), participants needed to phase-shift their real hand movements—and more strongly so depending on the amount of delay; in the RH task, no such real hand movement shift was required. We furthermore compared movement amplitudes between conditions, by determining the local hand flexion minima and maxima in each movement cycle, and comparing them across conditions. Since the hand movement data were not normally distributed (determined by the Shapiro test), we used the non-parametric Friedman’s test to assess statistical significance.

### fMRI data preprocessing and analysis

The fMRI data were recorded with a 3T scanner (MAGNETOM Prisma; Siemens) equipped with a 32-channel head coil. T2*-weighted images were acquired using an echo-planar imaging (EPI) sequence (voxel size = 3 mm^3^, matrix size = 64 x 64, 48 slices, TR = 2.36 s). Additionally, for each participant we recorded a T1-weighted “structural” image (voxel size = 0.85 mm^3^, matrix size = 320 x 320 mm, 240 slices, TR = 2.4 s) and a field map (voxel size = 3 mm^3^, matrix size = 64 x 64, 48 layers, TR = 532 ms, 48 layers). All fMRI preprocessing steps and analyses were performed using MATLAB (MathWorks) and SPM12 (Wellcome Trust Centre for Neuroimaging, University College London; https://www.fil.ion.ucl.ac.uk/spm). The functional images were realigned and unwarped using voxel displacement maps calculated from the field maps. These images were then spatially normalized to MNI space, resliced to 2 mm³ voxels, and smoothed with an 8 mm full width at half maximum Gaussian kernel. The T1-weighted structural images were also normalized (but not smoothed) and averaged to create a group averaged structural image (based on 19 participants, as one participants’ structural image was, unfortunately, lost). For each participant, we fitted a general linear model (GLM, 192 s high-pass filter) to the preprocessed data. The main purpose of our design was to identify potential task set dependent differences in the processing of delays and their changes; furthermore, we also looked at tracking error processing. Thus, we modelled each condition (VH task, RH task) as a continuous train of 1 s onset regressors (corresponding to a continuous 188 s block regressor) with three orthogonalized and mean-centered parametric modulators: The first parametric modulator modelled the amount of delay present during the current train of movements (see Fig. 1C); i.e., larger values corresponded to larger delays. The second parametric modulator modelled the time points at which the delay changed, weighted by the absolute amount of change; i.e., a larger change was assigned a larger value. Note that the delay and change modulators were, in each of the 6 runs, identical for the VH and RH task. The third parametric modulator reflected the modality specific hand-target tracking error; i.e., the mismatch between the virtual hand and the target in the VH task, and between the real hand and the target in the RH task. The values for the tracking error were calculated, for each grasping movement cycle (2s), as the average mean squared difference between the normalized trajectories of the target oscillation and the respective instructed (virtual or real) hand’s movement. As the parametric modulators were orthogonalized with respect to the condition regressor, they explained the variance associated with each manipulation around the respective condition’s mean. Furthermore, as the parametric modulators were orthogonalized with respect to each other, in the above order, the variance explained by each successive parametric modulator (regressor) can be interpreted as a unique effect over and above that explained by the previous modulators. Finally, we added the six realignment parameters as noise regressors to each run, alongside a session constant, and a regressor of no interest modelling the instruction periods.

On the first (participant) level, T-contrast images were created for each condition regressor vs baseline and for each parametric modulator, across all 6 runs per participant. These first-level contrast images were entered into group-level GLMs using paired t-tests with the factor ‘Task (VH, RH)’ for each regressor; in these GLMs, the second-level (group) T-contrasts were calculated. Activations obtained from the group-level contrasts were assessed for statistical significance using a voxel-wise threshold of *p*<0.05, family-wise error corrected (*p_FWE_*<0.05) for multiple comparisons. Our focus was on detecting blood oxygenation level dependent (BOLD) signal differences in brain regions processing visuomotor task parameters (including delays and their changes) *per se*. Therefore, we adopted a two-step procedure in our analysis: We first identified the brain regions responding to the main experimental manipulations *per se*; i.e., voxels showing significant main effects (i.e., positive correlations with the task, delay, delay change, and error regressors), thus defining our regions of interest. Then, we tested whether these regions would show significant between-condition (i.e., VH vs RH) differences; i.e., we evaluated and corrected the results of the differential contrasts within the search space of all significant voxels obtained from each respective main effect. For instance, we tested for significant BOLD signal increases by the VH > RH task within a search mask defined by significant (*p_FWE_*<0.05) activations by the main effect of task ((VH+RH) > rest). We have marked those significant effects with an asterisk in Table 2, for clarity. For completeness, we also tested for whole-brain significances in each contrast and for negative correlations with the regressors, which we report in the Supplementary material. The significant voxels resulting from the group level contrasts are presented as renders on SPM12’s template brains or superimposed onto the group averaged normalized structural image. The displayed significant activation maps contain only voxels surviving the chosen threshold of *p_FWE_*<0.05. For anatomical reference, we used the SPM Anatomy toolbox (Eickhoff et al., 2005) where possible.

### Functional connectivity analysis

Following the identification of significant differences in the delay-dependent BOLD signal between VH and RH tasks in the left cerebellum in our main analysis, we tested for delay-dependent changes in its functional connectivity to other brain areas. For this, we used psychophysiological interaction (PPI) analysis as implemented in SPM (Friston et al., 1997): We concatenated the experimental runs including all regressors for each participant, and from these first-level GLMs we extracted the cerebellar BOLD signal time course as summarized by the first Eigenvariate of all voxels within a 4 mm radius sphere centered on the participant specific peak effect of the delay (VH > RH) contrast, within 10 mm of the cerebellar group-level peak. The mean coordinates and standard deviations the cerebellar seeds were: x = -37.1 ± 4.0, y = -45.9 ± 4.2, z = -33.8 ± 5.0. We then calculated the interaction between the different effect of the parametric delay regressors of the VH > RH tasks (the “psychological variable”) and the extracted cerebellar BOLD signal time course (the “physiological variable”). To look for significant connectivity changes, the first-level contrast images calculated on this interaction effect were entered into a one-sample t-test on the group-level. In addition to a default whole-brain thresholded analysis, we specifically looked for connectivity changes in region of interests defined by those brain regions that showed an increased BOLD signal during the VH > RH task. These results were evaluated for significance using a voxel-wise statistical threshold of *p_FWE_*<0.05, within the significant voxels identified by the contrast “Task (VH > RH)”, see Fig. 3B. We did this because we expected the cerebellum, based on its well-known importance in visuomotor adaptation (Tzvi et al., 2022), to communicate specifically with brain regions engaged more strongly by the VH task (i.e., the task requiring such adaptation).

## Results

### Behavioral results

We first tested for task compliance; i.e., whether participants shifted their real hand movements more strongly in the VH task than in the RH task (as this shift of the real hand movements was required to compensate for the visuomotor delays in the VH task; i.e., to align the lagging visual movements with the target oscillation). As instructed, participants systematically shifted their real hand movements in the VH task (Fig. 2). Importantly, participants shifted their real hand movements generally more with increasing delays (see Table 1). This suggested a delay-dependent compensatory response, albeit partial (i.e., participants did not completely compensate for the delays; this was not surprising since we kept the individual delay periods relatively short in order to maximize the number of delay changes per run, see Materials and methods). Conversely, also as instructed, participant exhibited no such delay-dependent shift in the RH task, where their real hand movements were consistently lagging the target oscillation slightly. The overall difference in real hand shift between VH and RH task was significant (Friedman’s test, *χ²*(1)=83.23, *p*<.001, see Table 1 for means and standard deviations).

**Figure 2:**
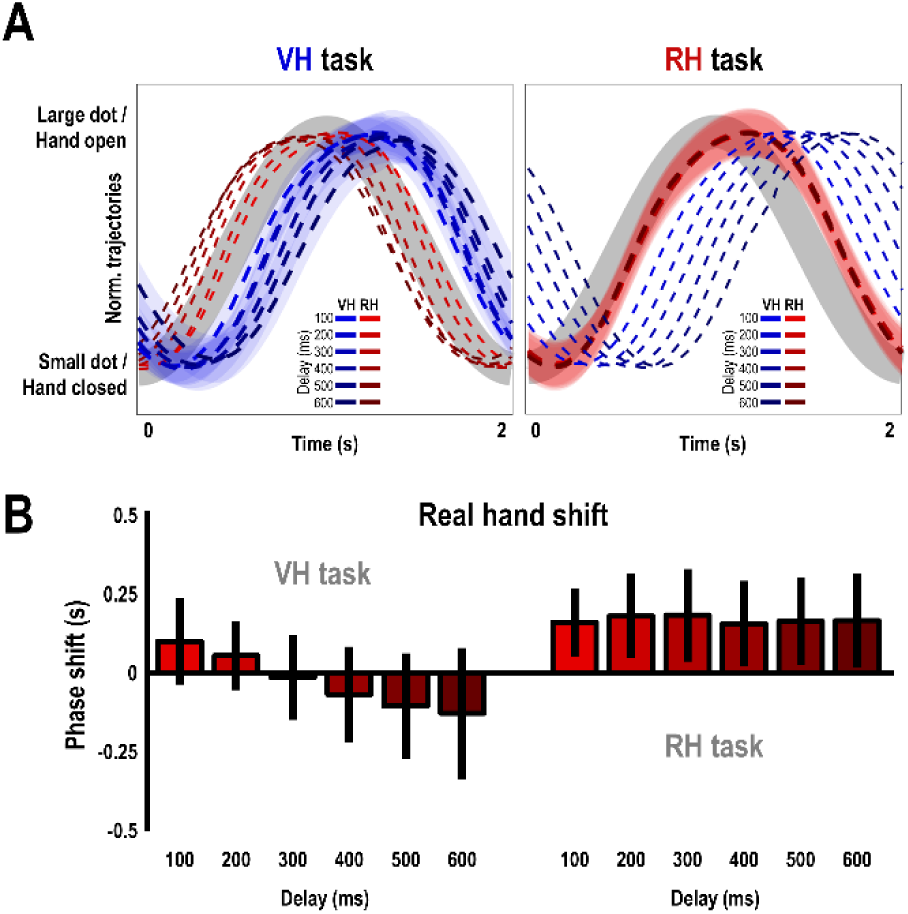
Behavioral performance. **A:** Hand-target tracking in the two tasks. The red and blue curves show the averaged normalized grasping trajectories of the real and virtual hand, relative to one cycle of the target’s 0.5 Hz oscillation (in grey), for of each delay level (excluding the first two cycles due to the gradual delay changes, see Methods). For display purposes, the respective instructed modality is plotted in bold, with shaded areas representing the associated standard errors of the mean. **B:** Average phase shifts of the real hand (in s with standard deviations) from the target oscillation per delay level in each task, indicating whether real hand movements were leading (negative values) or lagging (positive values) the target oscillation. Importantly, in the VH task (left plot), participants shifted their real hand movements ahead in time, whereby the shift increased proportionally with the amount of delay. Thus, participants exhibited the desired delay-dependent compensatory responses necessary to counteract the visuomotor delays and re-align the virtual hand with the target oscillation. Conversely, in the RH task (right plot), they kept their real hand movements equal across the different delay levels (i.e., successfully ignoring incongruent vision). The difference in real hand behavior between the VH and RH task was significant at *p*<0.001. See Table 1 for details.

**Figure 3:**
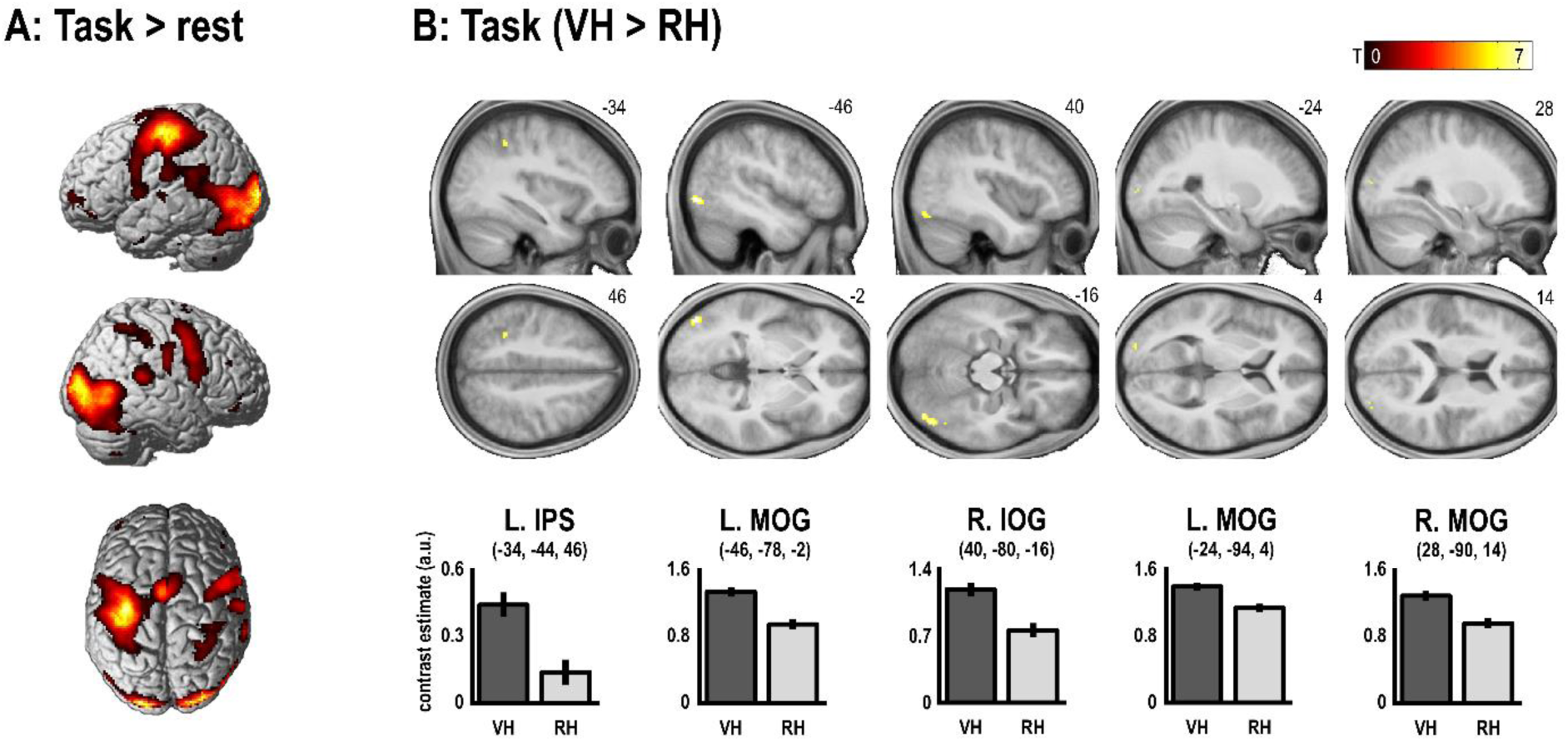
Brain activations related to hand-target tracking. **A:** Voxels showing a significant (*pFWE*<0.05) main effect of task > rest. **B:** Significantly (*pFWE*<0.05) stronger activations during the VH task > RH task were observed in the left IPS and in in the bilateral extrastriate visual cortices. No significant converse effects (RH > VH) were found. The bar plots show the contrast estimates of the VH and RH condition regressors from each peak voxel, with associated 90% confidence intervals. See Table 2 for details.

**Table 1.** Real hand behavior (means ± standard deviations).

| **Real-hand target phase shift (s)** |  |  |
| --- | --- | --- |
| *Delay (ms)* | *VH task* | *RH task* |
| 100 | 0.10±0.14 | 0.17±0.13 |
| 200 | 0.06±0.11 | 0.16±0.11 |
| 300 | -0.01±0.14 | 0.18±0.13 |
| 400 | -0.07±0.15 | 0.16±0.13 |
| 500 | -0.10±0.17 | 0.16±0.13 |
| 600 | -0.13±0.21 | 0.17±0.15 |
| *All* | -0.03±0.17 | 0.17±0.13 |

| **Movement amplitude (a.u.)** |  |  |
| --- | --- | --- |
| *Delay (ms)* | *VH task* | *RH task* |
| 100 | 0.46±0.10 | 0.47±0.09 |
| 200 | 0.47±0.09 | 0.48±0.09 |
| 300 | 0.46±0.09 | 0.48±0.09 |
| 400 | 0.45±0.10 | 0.49±0.10 |
| 500 | 0.47±0.10 | 0.50±0.09 |
| 600 | 0.45±0.09 | 0.47±0.10 |
| *All* | 0.46±0.09 | 0.48±0.09 |

Excl. first 2 movement cycles (see Materials and methods).

**Table 2.**
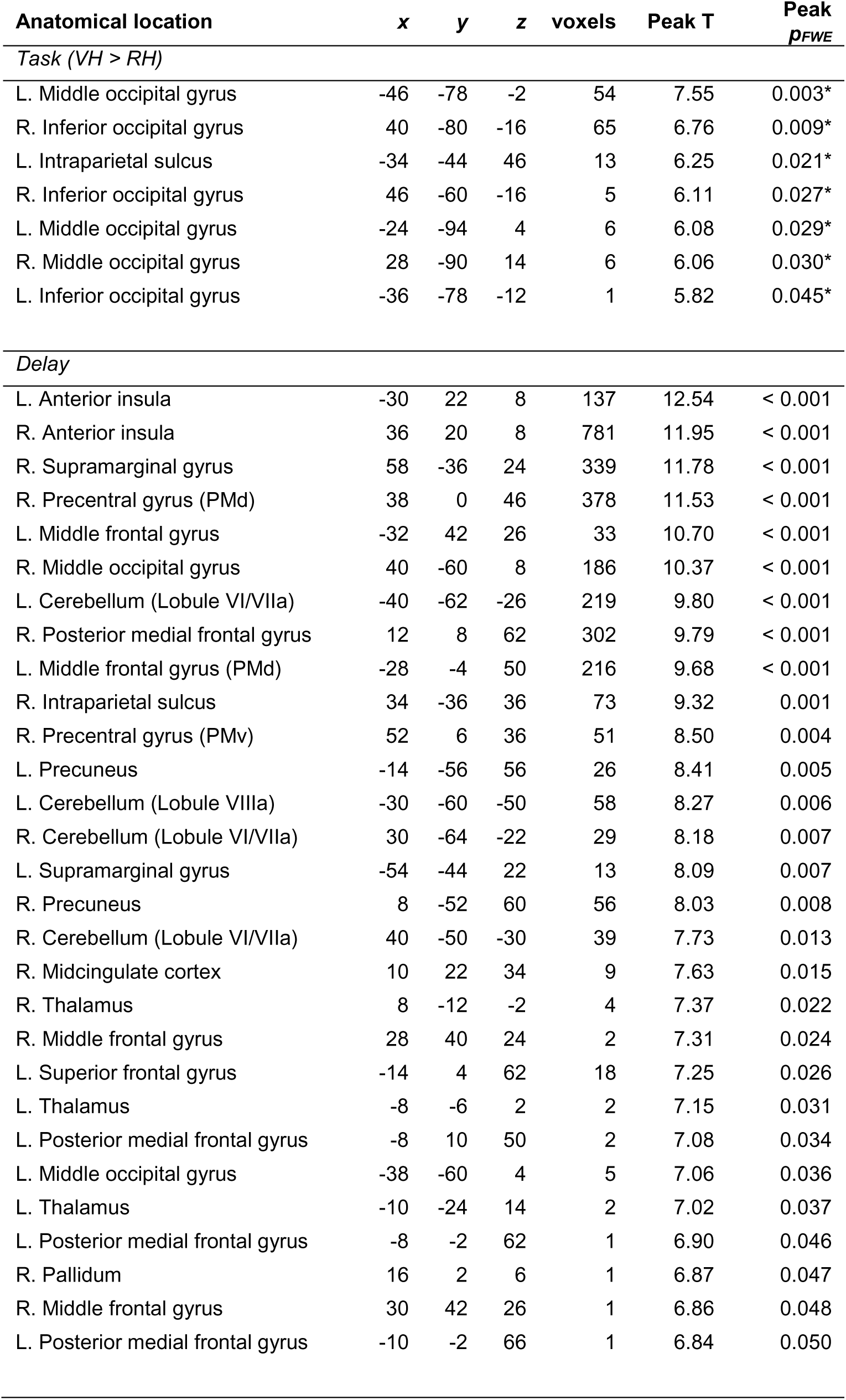
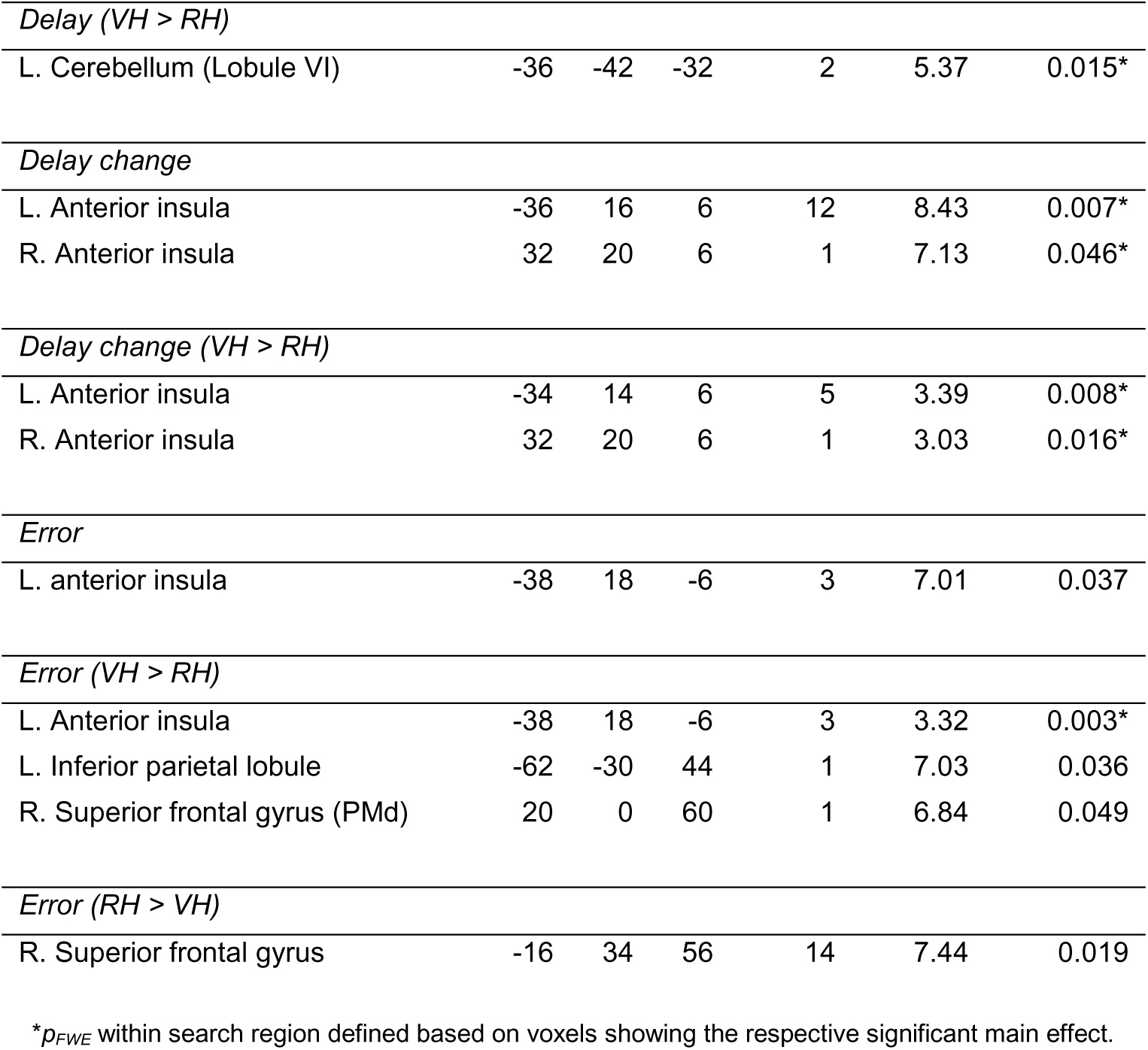
Significant activations obtained from the group-level contrasts.

On average, participants’ grasping movements were of somewhat smaller amplitude in the VH than in the RH task (Friedman’s test, *χ²*(1)=4.17, *p*<.05); however, this difference amounted to only ∼5% of the average amplitudes (Table 1). There was no significant effect of delay on movement amplitudes (Friedman’s test, *χ²*(5)=1.85, *p*=.87). In the first movement cycle (where the delay changes were introduced), participants made larger amplitude movements than in the remaining ones (Friedman’s test, *χ²*(1)=5.18, *p*<.05); again, with somewhat smaller amplitude in the VH than the RH task (Friedman’s test, *χ²*(1)=4.47, *p*<.05, Table S1). There was no significant effect of delay on movement amplitudes of the first movement cycle alone (Friedman’s test, *χ²*(5)=2.41, *p*=.79). Together, our behavioral results suggested good task compliance; i.e., that participants exhibited delay-dependent visuomotor compensation in the VH but not the RH task.

### fMRI results

In our fMRI analysis, we first looked for significant activations in the whole brain by hand-target tracking in both conditions (i.e., the main effect of task > rest). Significant (*p_FWE_*<0.05) voxels obtained from this contrast were located in the expected motor and visuomotor areas; including the left primary motor cortex, pre- and supplementary motor areas, the PPC, and visual areas (Fig. 3A). Within a search region defined by these task related activations, we found significantly (*p_FWE_*<0.05) stronger BOLD signal increases during the VH > RH task in the left IPS and in the bilateral extrastriate visual cortex, including middle and inferior occipital gyri (MOG/IOG, see Figure 3B and Table 2. At a more liberal, whole-brain threshold (*p*<0.001, uncorrected), further activation differences were observed in the bilateral posterior parietal, premotor, anterior insular cortices, and in the bilateral cerebellum (Fig. S1). The converse contrast, RH task > VH task, yielded no significant activation differences.

Next, we looked for brain regions increasing their responses proportionally with the amount of visuomotor delay; i.e., for voxels that would show a positive BOLD signal correlation with the parametric delay regressor. We found a significant (*p_FWE_*<0.05) main effect of delay in a wide-spread network of brain regions previously associated with visuomotor delay processing; with major clusters in the (predominantly right) temporoparietal and premotor cortex, the bilateral anterior insula, and in the bilateral cerebellum (Fig. 4A). Conversely, delay was associated with relatively suppressed BOLD signal in the right rolandic operculum (Fig. S2 and Table S2). There was no overlap between regions showing a significant main effect of delay and those showing a significant preference for the VH > RH task (no common voxel between Fig. 4A and Fig. 3B), but some delay-sensitive voxels in the premotor cortex and the left cerebellum showed a VH > RH task effect at a lower statistical threshold (*p* < 0.001, see Fig. S3).

**Figure 4:**
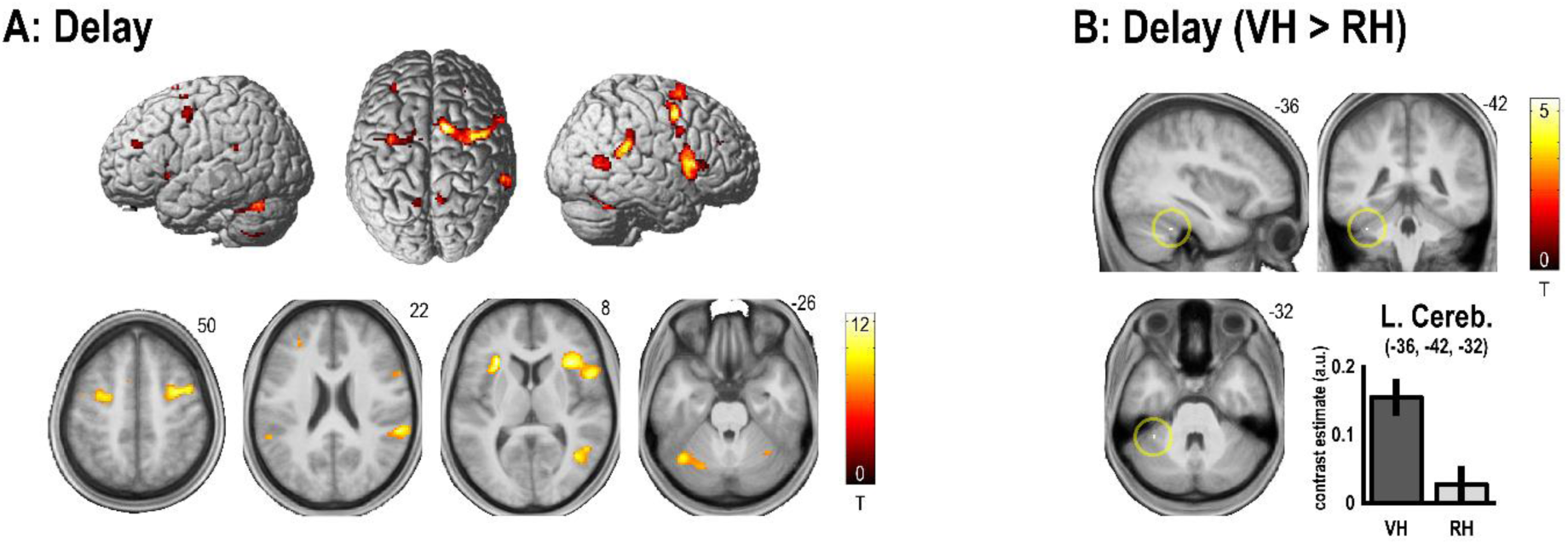
Significant BOLD signal correlations with visuomotor delay. **A:** Render and slice overlays showing significant (*pFWE*<0.05) BOLD signal correlations with the parametric delay regressor in premotor, insular, temporoparietal, and cerebellar regions. **B:** Significantly (*pFWE*<0.05) stronger delay-related BOLD signal increases during the VH > RH task were observed in the left anterior cerebellum (Lobule VI). The bar plots show the contrast estimates of the parametric delay regressor, with associated 90% confidence intervals, from the cerebellar peak voxel. See Table 2 for details.

Within the regions responding to delay, we found a significantly stronger delay-related BOLD signal increase during the VH > RH task in the left anterior cerebellum (Lobule VI, see Fig. 4B). No other significant correlation differences with delay were found (see Fig. S4). In voxels of the right temporoparietal cortex showing a main effect; i.e., the SMG and pSTS, delay related activity was slightly, but non-significantly stronger in the RH > VH task, see Fig. S5.

Together, these results mean that of all regions responding to visuomotor delays, only the cerebellum processed them significantly more strongly when the delay was task relevant; i.e., when participants were tracking the target with the virtual hand (trying to adapt to delays).

Following this result, we tested for related differences in the delay-dependent functional coupling of the left cerebellum between the VH and RH task. A PPI analysis revealed that during the VH > RH task, larger visuomotor delays were associated with a significantly increased coupling of the left cerebellum with regions in the right IPL (*x* = 58, *y* = -38, *z* = 48, *p_FWE_*<0.05), and with the left IPS (*x* = -34, *y* = -44, *z* = 46) and the right MOG (*x* = 48, *y* = -70, *z* = -8, *p_FWE_*<0.05, within a search mask defined by the contrast *Task (VH > RH)*). See Figure 5 (and Fig. S6 for an uncorrected render of the PPI results).

**Figure 5:**
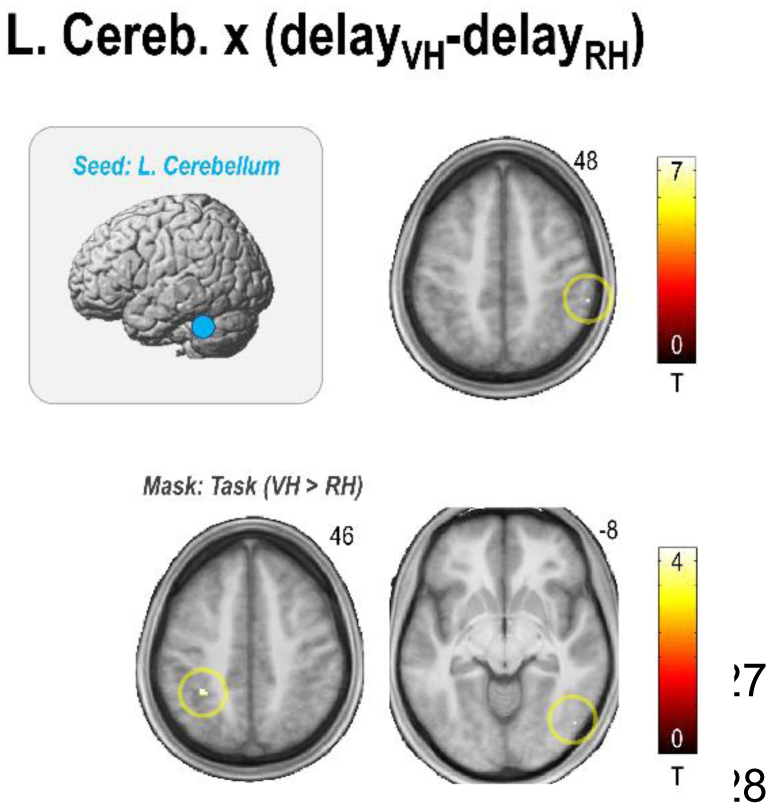
PPI analysis showing brain regions with stronger delay-dependent functional coupling with the left cerebellum in the VH > RH task. During the VH > RH task, visuomotor delay had a significantly (*pFWE*<0.05) stronger impact on the functional coupling of the left cerebellum with the right IPL, and with two of the brain regions showing a significant effect of VH > RH task: the left IPS and the right MOG.

Next, we searched for effects related to *changes* in visuomotor delay; i.e., for voxels showing a correlation with the parametric regressor encoding changes (transitions) in visuomotor delays— the “oddballs” in the roving design.

We found a significant (*p_FWE_*<0.05) main effect of delay change in the bilateral dorsal parts of the AI (Fig. 6A). At lower statistical thresholds, further activations emerged in the premotor and extrastriate visual cortex (Fig. S7). The AI thereby showed significantly stronger activation by delay changes during the VH task than the RH task (*p_FWE_*<0.05, within voxels showing a main effect, see Fig. 6B). Delay changes were, conversely, associated with relatively suppressed BOLD signal in the left S1 (Fig. S8 and Table S2). No other significant differences in delay change related activations between the VH and RH task were found.

**Figure 6:**
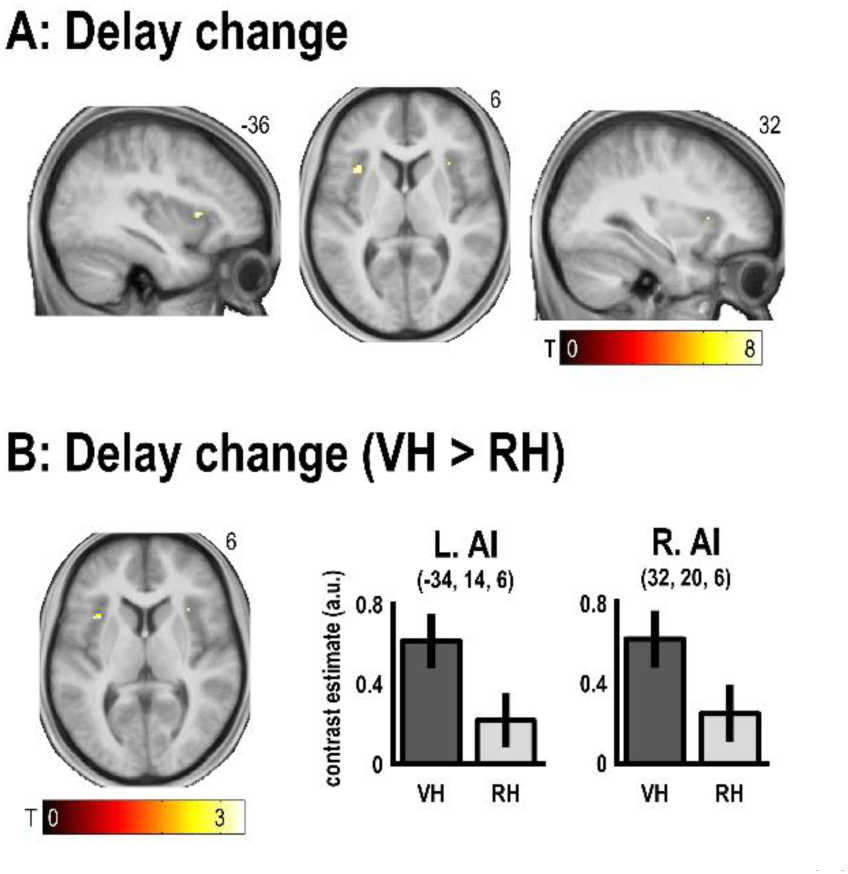
Significant activations by delay changes (“oddballs”). **A:** The bilateral dorsal AI was significantly activated by changes in visuomotor delay (*pFWE*<0.05). **B:** Delay changes correlated with the BOLD signal in the left and right AI significantly more strongly in the VH task > the RH task. The bar plots show the contrast estimates of the parametric delay change regressor from each peak voxel, with associated 90% confidence intervals.

Finally, although this was not the main aim of our design, we looked for brain activations related to hand-target tracking error (i.e., correlations with the deviation of the virtual hand’s grasping movements from the target in the VH task, and of the unseen real hand’s grasping movements from the target in the RH task). Tracking error significantly correlated with activity in the left ventral AI (*p_FWE_*<0.05, see Fig. 7A). No other significant (whole-brain) activations were found; further notable activations that did not survive correction for multiple comparisons (*p*<0.001) were in the supplementary motor area and the left PPC (see Fig. S9). Conversely, a whole-brain analysis revealed significant inverse BOLD signal correlations with error in the left M1, and in visual and temporal areas (Fig. S9 and Table S2). In the left AI, BOLD correlated significantly more strongly with tracking error during the VH task than during the RH task (Fig. 7B). A whole-brain analysis revealed further significantly (*p_FWE_*<0.05) stronger BOLD signal correlations with tracking error during the VH task > the RH task in the left IPL and in the right PMd (Fig. S10). The converse contrast (error RH > VH) revealed significant voxels in the left superior frontal gyrus (Fig. S9).

**Figure 7:**
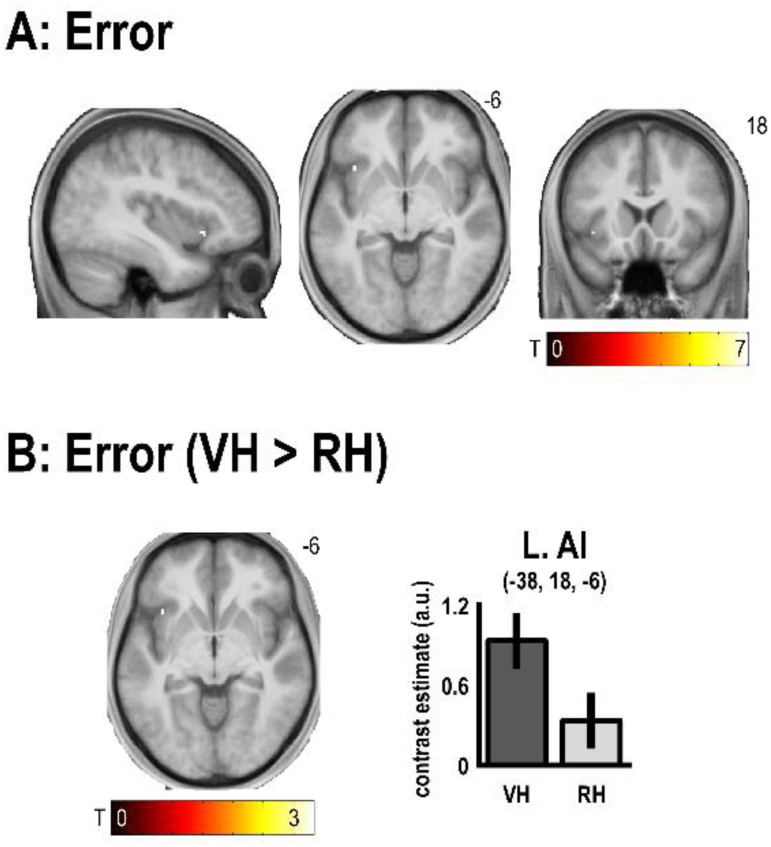
Significant correlations with hand-target tracking error. **A:** The left ventral AI showed a significant BOLD signal correlation with hand-target tracking errors (*pFWE*<0.05). **B:** Hand-target tracking errors correlated significantly more strongly with left AI BOLD in the VH > RH task. The bar plots show the contrast estimates of the parametric error regressor with associated 90% confidence intervals. See Table 2 for details.

## Discussion

Using a virtual reality based, continuous, right hand-target tracking task with variable visuomotor delays, we were able to induce a behavioral focus onto or away from incongruent visual movement feedback. Participants were able to adapt to changing visual feedback delays by adjusting their real hand movements in the VH task, whereas they successfully kept their unseen real hand aligned with the target in the RH task; i.e., ignoring the visual feedback delays and their changes.

Our fMRI analysis revealed significant and anatomically distinct BOLD signal correlations with the different task parameters: Firstly, when participants had to use visual movement feedback for target tracking (i.e., tracking with the delayed virtual hand movements vs with the unseen real hand), we observed significant BOLD signal increases in the left (contralateral) IPS and in the bilateral extrastriate visual cortices. Note that virtual hand movements were always delayed, therefore, the VH condition effectively constituted a continuous visuomotor integration task. Thus, this result supports the well-documented importance of the PPC, specifically the IPS, for visually guided action control (Christensen et al., 2007; Della-Maggiore et al., 2004; Desmurget et al., 1999; Grefkes et al., 2004; Limanowski et al., 2017; Ogawa et al., 2006). Furthermore, our results speak to previous work associating activity in similar regions of the left PPC/IPS with planning and executing hand movements under incongruent visual movement feedback; suggesting visuomotor integration as a key function of the left PPC (Grafton et al., 2008; Kuang et al., 2015; Limanowski & Friston, 2020; Lindner et al., 2010; Quirmbach & Limanowski, 2024).

The extrastriate effects very likely reflect the stronger activation of occipitotemporal motion sensitive areas V5/MT by visual movements in the VH > RH task. Conveying visual movement information via the MT/LOC plays an important part in upper limb action planning and control (Lingnau & Downing, 2015). That cognitive factors such as attentional set can bias MT responses to visual motion has been demonstrated previously (Büchel, 1998; Maunsell & Treue, 2006). We previously found similar increased putative V5/MT activations by hand-target tracking when visual movement feedback was task relevant vs irrelevant (Limanowski & Friston, 2020). The present results therefore corroborate the idea of a contextually sensitive processing of visual movement feedback in the lateral extrastriate visual cortex. Although we can only speculate about this, the contextual control over these attentional effects could stem from the IPS (Kanwisher & Wojciulik, 2000). In sum, these activation differences can be interpreted as capturing the effects of an instructed cognitive-attentional task set (i.e., integrating delayed visual movement feedback with motor plans, or ignoring it in favor of proprioceptive movement feedback).

Second, over and above the task set effects, our parametric delay regressor captured effects specifically related to the amount of visuomotor mismatch present during movement. We found significant BOLD signal correlations with the delay regressor across both VH and RH tasks in a set of temporoparietal, premotor, posterior parietal, and cerebellar brain regions; with the strongest and most extensive effects in the bilateral AI, the right SMG, pSTS, and premotor cortex, and the bilateral cerebellum. This anatomical pattern replicates the results of many previous studies, in which visuomotor incongruence-related BOLD signal increases were observed in similar brain regions, frequently also with right-hemispheric dominance (Balslev, 2004; Leube, 2003; Limanowski et al., 2017; Nahab et al., 2011; Ogawa et al., 2007; Ohata et al., 2020; Tsakiris et al., 2010; Van Kemenade et al., 2017, 2019). The most common interpretation of such incongruence (e.g. delay) dependent activity increases is that these regions act as ‘comparator’ modules that register mismatches between sensations predicted by the motor system and the actual sensory feedback (cf. Farrer & Frith, 2002; Balslev, 2004; Leube, 2003; Limanowski et al., 2018; Van Kemenade et al., 2017, 2019). Generally, our delay main effect supports this idea. But these activations could also relate to attentional reorienting or supervisory control (Cieslik et al., 2015; Corbetta et al., 2008) or, in adaptation tasks specifically, to feedback learning, action reprogramming, and sensorimotor adaptation (Grafton et al., 2008; Hartwigsen et al., 2012; Tzvi et al., 2022). With our design, we were able to test for differences in delay processing in the above areas between tasks involving visuomotor adaptation (VH) vs no adaptation (RH).

Crucially, only Lobule VI of the left anterior cerebellum showed a significantly stronger correlation with the amount of delay in the VH > RH task (the correlation was practically nonexistent in the RH task). In previous works, cerebellar activity has been associated with visuomotor adaptation (Block & Celnik, 2013; Chapman et al., 2010; Danckert et al., 2008; Küper et al., 2014; Luauté et al., 2009; Tzvi et al., 2020), but, in other works, also with the conscious detection of visuomotor delays (Arikan et al., 2019; Leube, 2003). In our study, the cerebellar delay dependent response difference between the VH and RH task was not simply related to the mere detection of visuomotor mismatches (because these were identical in the VH and RH tasks), nor to signaling unexpected stimulus combinations (i.e., there was no significant effect of delay *change* between VH and RH tasks in the cerebellum). Neither can the cerebellar responses be explained by different effects of visual vs proprioceptive hand-target tracking per se (cf. the VH > RH task effect observable in the IPS and LOC). Therefore, we suggest that the observed cerebellar responses related specifically to visuomotor adaptation; i.e., the repeated adaptation of forward models by unpredicted visual movement feedback, when vision was needed for target tracking. This interpretation speaks to the notion of cerebellar forward models of sensory action consequences, which can be updated by sensory prediction errors to enable adaptation (Miall et al., 1993; Shadmehr et al., 2010; Tseng et al., 2007; Synofzik et al., 2008; Wolpert, Miall, et al., 1998).

Moreover, cerebellar connectivity to the PPC—the right IPL, and furthermore, also the left IPS region showing a VH > RH task effect—was more strongly correlated with the amount of visuomotor delay in the VH > RH task. In other words, when the tracking task required repeated adaptation to delayed visual movement feedback (i.e., in the VH but not the RH task), the cerebellum not only increased activity under larger delays more strongly, but also increased its connectivity to the PPC more strongly. We believe the increased cerebellar coupling with the PPC (a region more strongly involved in the VH task per se, see above) could indicate the communication of the adjusted forward model output for state estimation and visuomotor tracking coordination in the PPC (cf. Christensen et al., 2007; Miall et al., 2001). Thus, our results support the idea of flexible forward models for visuomotor planning and adaptation in the cerebellar-posterior parietal network (Chapman et al., 2010; Kuang et al., 2015; Miall, 2003; Quirmbach & Limanowski, 2024; Tzvi et al., 2022).

Interestingly, the differential delay effect was significant in the left cerebellar hemisphere (i.e., contralaterally, but also detectable ipsilaterally at *p*<0.001). While one would generally expect cerebellar motor-related activity more strongly ipsilaterally to the moving hand, motor learning seems to activate the cerebellum bilaterally (Hardwick et al., 2013). Notably, some studies, using right-hand responses as well, have associated delay detection and adaptation, and agency evaluation specifically with the left cerebellum as well (Arikan et al., 2019; Kilteni & Ehrsson, 2024; Kufer et al., 2024; Yomogida et al., 2010). In this light, one could speculate about a potential similarity with studies implicating the left cerebellum (and left frontoparietal cortical regions) with temporal attention and orienting (Coull & Nobre, 1998; Ivry et al., 2002; Schubotz et al., 2000). This links to the outstanding question about the potential difference between the processing and adaptation to temporal vs spatial visuomotor incongruencies, and whether those recruit different brain regions (cf. Rohde & Ernst, 2016; Vigh & Limanowski, 2025). This should be addressed by future work.

In notable contrast, we found no significantly different BOLD signal responses to visuomotor delay during the VH vs RH task in or around the temporoparietal junction—regions previously implied in the comparison of predicted and actual visual movement feedback (see Introduction)—despite a very strong correlation with delay in both tasks (i.e., main effect). Furthermore, the delay-sensitive temporoparietal regions (i.e., the SMG and pSTS) did not show any significantly different activation during the VH vs RH tasks *per se*. This means that in these regions, unlike in extrastriate visual cortex, the instructed (attentional) task set did not influence visual feedback (delay) processing. Similarly, visual feedback changes were not processed differently depending on task instruction (unlike in the AI, see below). Thus, in our experiment, the temporoparietal cortex—among other delay sensitive regions—processed visual movement feedback and visuomotor delays similarly, i.e. irrespective of whether visual feedback was task relevant and adapted to, or task irrelevant and ignored. Therefore, our results support the idea of the temporoparietal cortex acting as a general ‘comparator’ of predicted and actual visual movement feedback (Balslev, 2004; Brass et al., 2009; Farrer et al., 2003, 2008; Jeannerod, 2004; Ohata et al., 2020; Tsakiris et al., 2010; Uhlmann et al., 2020; Van Kemenade et al., 2017, 2019). Our results now show that the temporoparietal comparison is based on a fundamental ‘background’ process—presumably, of self-other distinction and action attribution—that is *not* influenced by the behavioral relevance of visual feedback.

It should be noted, however, that upon closer inspection, the delay-sensitive temporoparietal regions (i.e., the SMG and pSTS) showed slightly, non-significantly stronger delay related responses in the RH task than in the VH task. Tentatively, this could suggest an effect of visuomotor adaptation; i.e., as the cerebellar forward predictions of visual movement consequences were slowly adapted to the delays in the VH task (but not the RH task), the delayed virtual hand movements would now constitute less of a mismatch to the temporoparietal comparators. However, as our experiment was not designed with full adaptation and its temporal evolution mind (see Materials and methods), this speculation has to be evaluated by future work. Thirdly, we observed significant a main effect of changes in delay in the bilateral AI, while the left AI also showed a significant main effect of hand-target tracking errors. These results replicated our previous findings obtained in similar hand-target phase matching tasks (Limanowski et al., 2017; Quirmbach & Limanowski, 2022). They also align with the well-established role of the AI in error detection and performance monitoring (Pereira et al., 2023; Ullsperger et al., 2010). Furthermore, they could suggest a preferential processing of unpredicted visual movement feedback, in line with previous fMRI studies linking AI activity to visuomotor learning and control (Grafton et al., 2008; Quirmbach & Limanowski, 2022). Previous studies have also indicated similar, comparable neural responses to novel events and errors in the AI (among other regions, (Notebaert et al., 2009; Ullsperger et al., 2014). Crucially, both AI BOLD signal correlations (with delay change and error) were significantly stronger when tracking the target with the virtual than the real hand. This suggests a specific function of the AI in continuous visuomotor control; namely, the processing of visual mismatch signals—including changes in visuomotor mappings and visually conveyed performance errors. Thus, our results support the idea of a task-dependent processing of ‘salient’ signals, such as behavioral errors or behaviorally relevant changes in task parameters, in the AI (Corbetta et al., 2008; Harsay et al., 2018; Higo et al., 2011; Sridharan et al., 2008; Ullsperger et al., 2010; Yomogida et al., 2010; Yucel et al., 2007). Potentially, the stronger AI activity in the visuomotor adaptation (VH) task could also be related to its role in action-outcome self-attribution (“agency”, cf. Farrer & Frith, 2002; Seghezzi et al., 2019; Sperduti et al., 2011), but we cannot draw any conclusions on this based on our design.

Our results need to be interpreted in light of the following limitations: Per design, visuomotor delays required (brief) adaptation in hand movements in the VH task, but constant hand movements in the RH task. The tasks thus likely differed in difficulty, which was visible in the overall better hand-target alignment in the RH than the VH task. Movement amplitudes were slightly but significantly different between tasks, which could have affected motor related activity. However, we observed no significant BOLD signal differences between the VH and RH task in primary motor or somatosensory areas. Furthermore, we ordered our parametric modulators to reflect first the experimental manipulations that were the focus of our experiment (i.e., delay and its changes), which might have limited the variance available for the error regressor to explain. This could explain why further common regions of the error-processing network (Ullsperger et al., 2014) were only detectable at uncorrected thresholds. Per our task design, errors may also have been more easily detectable in the VH task (through vision i.e. the virtual hand) than in the RH task (through proprioception i.e. the unseen real hand). Furthermore, we were unable to track eye movements inside the scanner; hence, we cannot rule out that some of the observed BOLD signal differences partly related to oculomotor behavior or eye-hand coordination. However, our non-spatial, central visual target rendered eye-movements unnecessary in this task; we have previously demonstrated that participants are well able to maintain central fixation in these kinds of tasks (Limanowski & Friston, 2020; Quirmbach & Limanowski, 2024).

To conclude, in a novel, continuous movement task involving visuomotor adaptation vs no-adaptation, we could distinguish between the roles of the PPC (visuomotor integration and visually guided action control), the cerebellum (forward model updates), temporoparietal regions (task-independent visuomotor comparisons), and the AI (signaling behaviorally relevant visuomotor changes) in the contextual control of movements under visuomotor conflict.

## Acknowledgments

This work was funded by the German Research Foundation (DFG, Deutsche Forschungsgemeinschaft) as part of Germany’s Excellence Strategy – EXC 2050/1 – Project ID 390696704 – Cluster of Excellence “Centre for Tactile Internet with Human-in-the-Loop” (CeTI) of Technische Universität Dresden. JL was supported by a Freigeist Fellowship of the VolkswagenStiftung (AZ 97-932).

## Data availability statement

The behavioral and fMRI data will be made available upon request. The data are not available publicly due to ethical and privacy restrictions.

