## Supplementary Material for "Task dependent cortico-cerebellar responses to delayed visual movement feedback"

Gesche Vigh[1,2] and Jakub Limanowski[2,3]
[1]Faculty of Psychology, Technical University of Dresden, 01069 Dresden, Germany
[2]Center for Tactile Internet with Human-in-the-Loop, Technical University of Dresden, 01187 Dresden, Germany
[3]Institute of Psychology, University of Greifswald, 17486 Greifswald, Germany

---

**Table S1.** Movement amplitudes in the first cycle (means ± standard deviations).

| **Movement amplitude (a.u.)** | | |
|---|---|---|
| *Delay (ms)* | *VH task* | *RH task* |
| 100 | 0.51±0.13 | 0.52±0.11 |
| 200 | 0.53±0.11 | 0.56±0.10 |
| 300 | 0.51±0.12 | 0.55±0.11 |
| 400 | 0.50±0.13 | 0.56±0.11 |
| 500 | 0.52±0.13 | 0.57±0.10 |
| 600 | 0.52±0.13 | 0.52±0.13 |
| *All* | *0.52±0.12* | *0.55±0.11* |

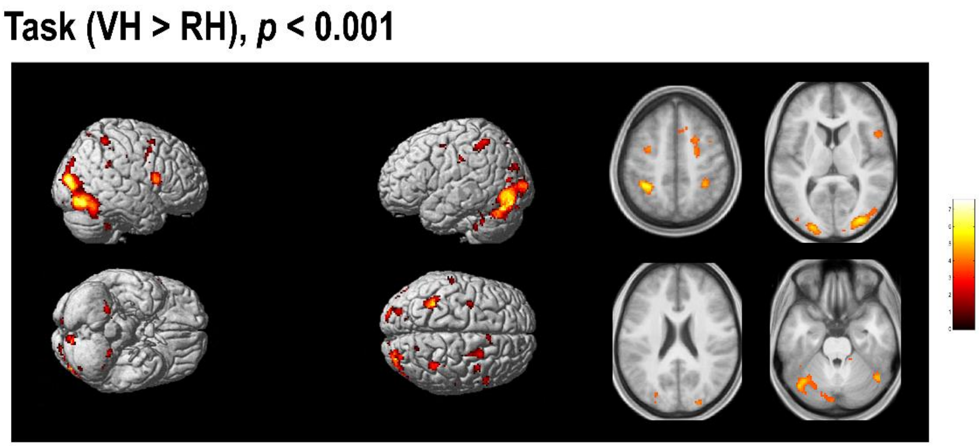


**Figure S1.** Uncorrected BOLD signal increases in the VH > RH task, related to Fig. 3B.

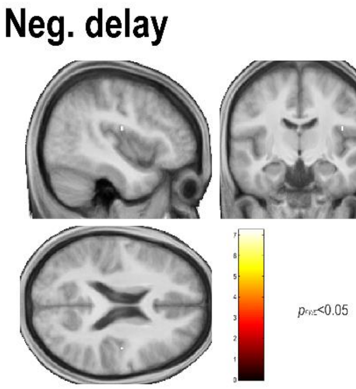


**Figure S2**. Significant negative correlations with delay in the right Rolandic operculum (area OP3). Related to Fig. 4.

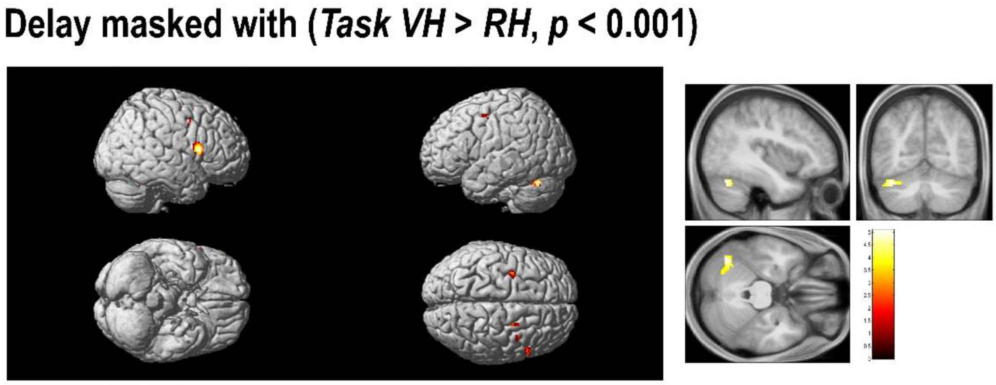


**Figure S3**. Delay sensitive voxels showing an effect of VH > RH task at *p*<0.001, uncorrected. Related to Figs. 3B and 4A.

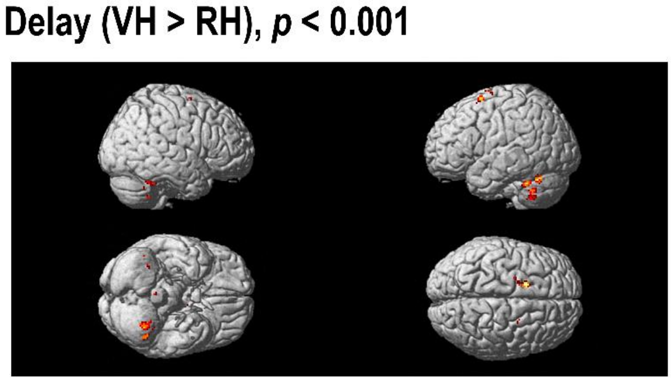


**Figure S4.** Uncorrected correlation differences for the contrast Delay (VH > RH), related to Fig. 4B.

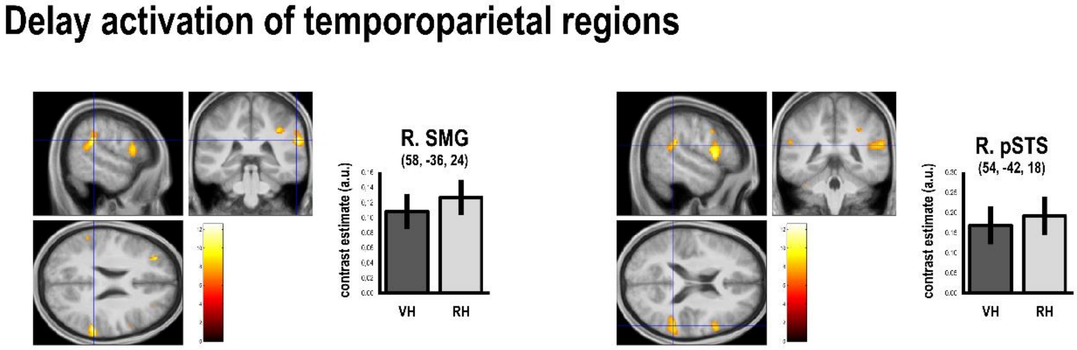


**Figure S5.** Delay-dependent activation of temporoparietal regions, related to Fig. 4A. The right SMG and pSTS both showed slightly, but non-significantly stronger activation by delays in the RH > VH task. The SPMs show the activation peaks from the main effect of delay.

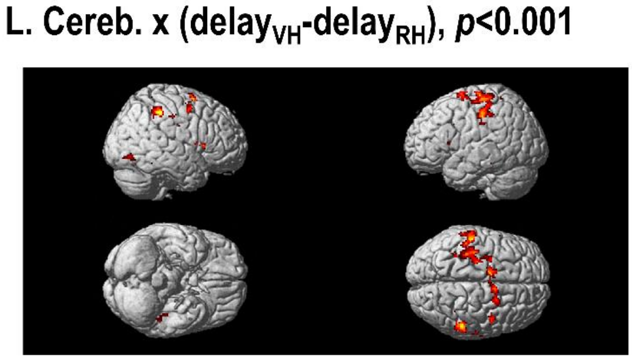


**Fig. S6**. Uncorrected results of the PPI testing for cerebellar connectivity, related to Fig. 5.

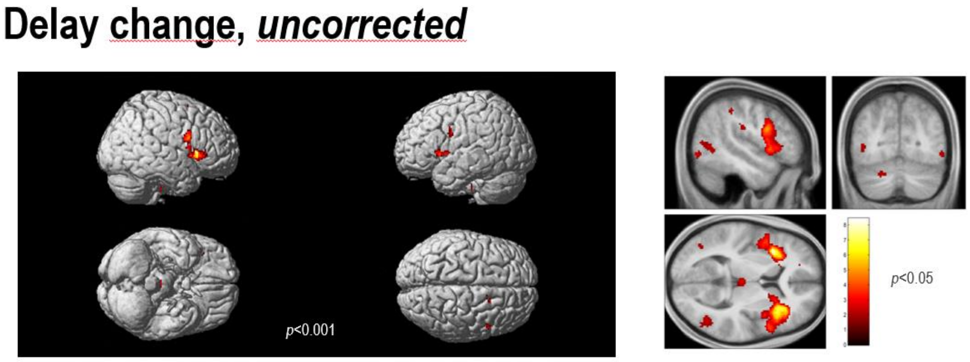


**Figure S7.** Uncorrected correlations with the delay change regressor, related to Figure 6. Extrastriate activations were observed at lower thresholds (right). It should be noted that in our previous study (Limanowski et al., 2017), changes in visuomotor mappings activated the bilateral extrastriate visual cortex much more strongly as in the present study. As these changes were between delayed and synchronous mappings (i.e., task periods requiring visuomotor adaptation or not), the extrastriate activations could perhaps be related to changes in (visual) attentional task set as captured by the task regressor in the present study.

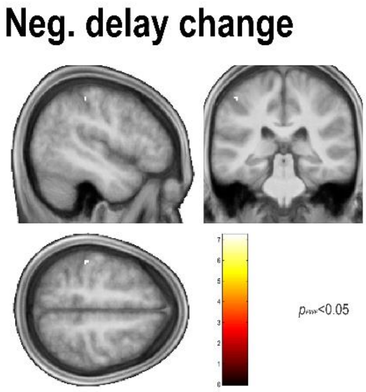


**Figure S8.** Significant negative correlations with delay change, see also Table S1. Related to Figure 6.

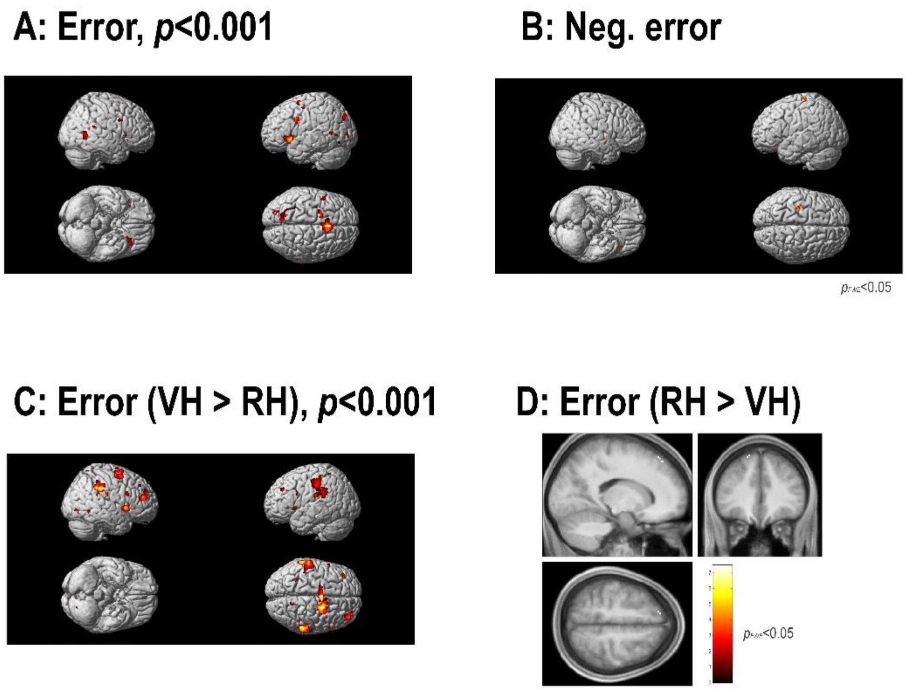


**Figure S9. A:** Uncorrected correlations with hand-target tracking error. **B:** Significant negative correlations with tracking error. **C:** Uncorrected correlation differences with error in the VH > RH task. **D:** Significant correlation difference with error in the RH > VH task. Related to Fig. 7.

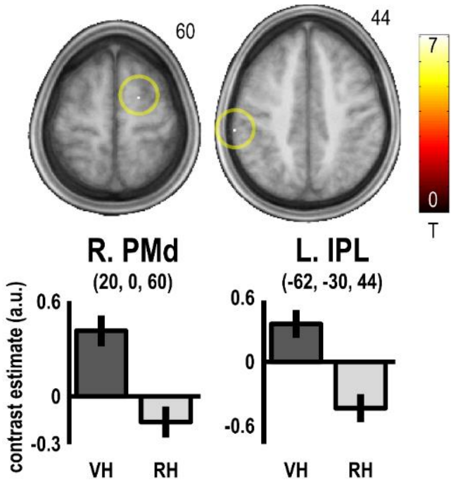


**Figure S10.** In a whole-brain analysis, the PMd and left IPL showed significantly ($p_{FWE}$<0.05) stronger correlation with tracking error in the VH > RH task. Related to Fig. 7.

**Table S2.** Significant voxels obtained from the negative group-level contrasts.

| Anatomical location | *x* | *y* | *z* | voxels | Peak T | Peak $p_{FWE}$ |
|---|---|---|---|---|---|---|
| *Neg. delay* | | | | | | |
| R. Parietal operculum (OP3) | 42 | -12 | 20 | 2 | 7.22 | 0.028 |
| *Neg. delay change* | | | | | | |
| L. Postcentral gyrus (BA2) | -46 | -32 | 54 | 4 | 7.23 | 0.039 |
| *Neg. error* | | | | | | |
| L. Precentral gyrus | -32 | -24 | 70 | 18 | 8.41 | 0.005 |
| R. Lingual gyrus | 10 | -76 | 0 | 13 | 7.86 | 0.010 |
| L. Inferior frontal gyrus | -40 | 26 | -18 | 3 | 7.63 | 0.014 |
| R. Superior temporal gyrus | 60 | -4 | -4 | 6 | 7.59 | 0.015 |
| L. Inferior frontal gyrus | -26 | 34 | -12 | 1 | 6.94 | 0.042 |
